# A smartphone-based genetically recombinant whole-cell biosensor for highly sensitive monitoring of polychlorinated biphenyls (PCBs)

**DOI:** 10.1101/2024.06.28.601110

**Authors:** Qiangqiang Luo, Faying Zhang, Mengjie Zhang, Shantong Hu, Xin Li, Li Pan, Zhenghui Lu, Pan Wu, Guimin Zhang

## Abstract

Polychlorinated biphenyls (PCBs) are highly carcinogenic and persistent pollutants commonly found in ecosystems. Their complex congeners pose a huge challenge to instrumental analysis and ELISA methods, which prefer single and known targets. To overcome this limitation, here we developed an *Escherichia coli* whole-cell biosensor (WCB) for simultaneously detecting multiple PCB congeners. In this sensor, PCBs were firstly converted into hydroxylated PCBs (OH-PCBs) by *bphAB* degradation circuits, which then serve as high-affinity targets of transcriptional factor HbpR_CBP6_-based sensing pathways for sensitive response through extensive chassis screening. The resulting biosensor BL21(DE3)/HbpR_CBP6_-*bphAB* shows the lowest detection limits for 2-CBP (2-chlorobiphenyl) to date and can recognize various PCB homologues, including 3-CBP, 4-CBP, 2,3-diCBP and 2,2’-diCBP, with detection limits of 0.06-1 μM. Further investigation of the docking structure and binding energy reveal that HbpR_CBP6_ has a stronger affinity for OH-PCBs than for PCBs, indicating that the conversion of PCB by BphAB enzymes is a key step to improve the sensitivity of WCB. Subsequently, we developed an immobilized hydrogel WCB and a smartphone-based detection procedure to facilitate real-time and user-friendly PCB detection. This study will not only advance the biomonitoring of PCB contaminants but also provide an innovative strategy for developing metabolic pathway-sensing proteins combined biosensor.

**Graphical Abstract:** 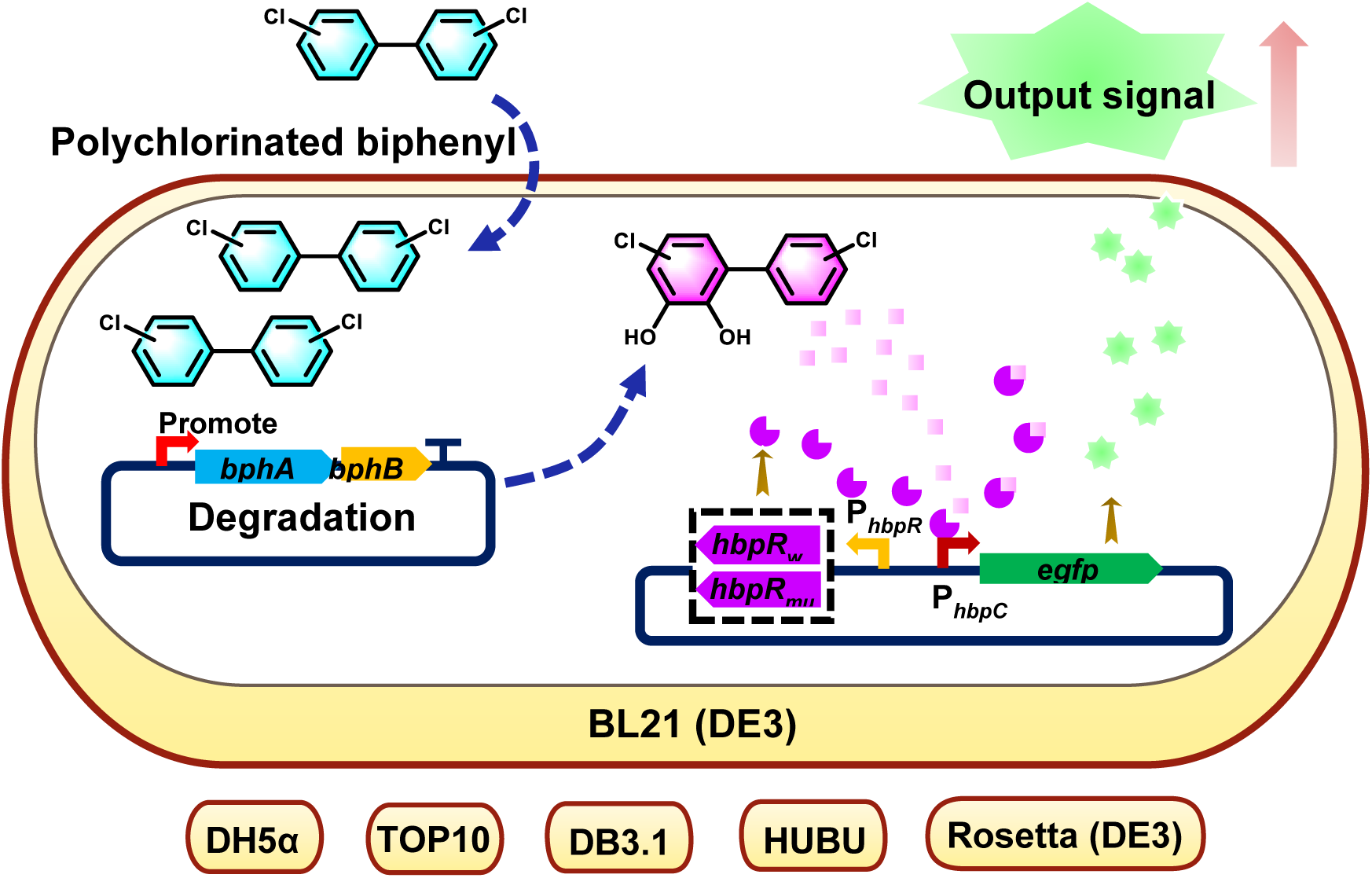

## 1. Introduction

Polychlorinated biphenyls (PCBs) are a class of chlorinated biphenyl compounds produced by replacing hydrogen atoms at different positions with chlorine atoms, including 2-chlorobiphenyl (2-CBP), 2,2’-dichlorinated biphenyl (2,2’-CBP), 2,4,4’-trichlorobiphenyl (2,4,4’-CBP), etc. PCBs were originally used for various industrial purposes, such as dielectric and heat transfer fluids, lubricants, flame retardants, and plasticizers (Erickson and Kaley, 2011). Due to their high persistence and toxicity, many countries have explicitly banned its use and it has been classified as a human carcinogen (Class I) by the International Agency for Research on Cancer (Lauby-Secretan et al., 2013). However, hundreds of millions of kilograms of PCBs are currently released into the environment(Zhang et al., 2021). The major threat to human health from PCBs in the environment is through bioaccumulation in the food chain (Qadeer et al., 2019; Rao et al., 2022; Richter et al., 1994). Therefore, monitoring PCBs is of global concern and effective techniques are urgently needed for detecting PCBs in routine samples (Wania and Mackay, 1993, 1996).

The diversity of environmental samples and the composition of PCB homologues pose great challenges to traditional instrument-based analytical techniques, such as gas chromatography-mass spectrometry (GC-MS) or high-performance liquid chromatography (HPLC) and ELISA methods that prefer to single and known targets (Deng et al., 2002; Erickson, 2018; Johnson et al., 2001; Reddy et al., 2019; Tsutsumi et al., 2008). In this context, whole-cell biosensors (WCBs) offer a promising alternative that not only alleviates the limitations of expensive instruments and expertise, but also enable simultaneous detection of multiple homologues (Bickman et al., 2018). Moreover, by integrating adsorption and metabolic circuits, WCBs can be extended to *in-situ* environmental bioremediation (Commault and Weld, 2016; Zhu et al., 2024). Nevertheless, due to the lack of direct sensing proteins, almost no WCBs have been reported yet for PCBs detection.

In fact, some researchers have tried to monitor PCBs indirectly by detecting PCB degradation intermediates. For example, through the upstream biphenyl pathway, PCBs are decomposed into chlorobenzoic acids (CBAs), which can be detected by WCBs containing *P_m_-gfp* sensing circuit. The upstream biphenyl pathway typically involves four enzymes: biphenyl 2,3-dioxygenase (BphA), cis-2,3-dihydro-2,3-dihydroxybiphenyl dehydrogenase (BphB), 2,3-dihydroxybiphenyl 1,2-dioxygenase (BphC) and 2-hydroxy-6-phenylhexa-2,4-dienoate hydrolase (BphD) (Pieper, 2005). Another example is that Liu’s research group combining the BphABCD degradation pathway with CBAs-induced *P_m_-gfp* sensing circuit to construct a *Pseudomonas fluorescens* F113L::1180-GFP to monitor degradation product of PCBs in soil (Liu et al., 2010). However, *P. fluorescens* produces natural yellow-green fluorescent pigment that interferes with the GFP reporter signal.

The rapid advancement in synthetic biology and genetic engineering has enabled the genetic identification of whole-cell biosensors by integrating heterologous metabolic pathways and transcriptional elements into designated chasses strains (Chen et al., 2023). For example, the HbpR/PhbpC sensing pathway from *P. azelaica* HBP strain was transplanted into an *E. coli* strain to prepare WCB for the detection of 2-hydroxybiphenyl (2-HBP) (Jaspers et al., 2001). Based on this, Siham et al. screened a variant of HbpR (I101V/D128N) and named it HbpR_CBP6_. The biosensor based on this variant displays a 20-fold increase in sensitivity to 2-HBP and an expanded response to 2-CBP, with a detection limit of 1.5 μM. However, this biosensor shows a weak or no response to homologous PCBs (Beggah et al., 2008). It seems that HbpR_CBP6_ has a higher affinity for hydroxylated biphenyl (OH-PCBs). We speculate that if PCBs can be converted to OH-PCBs, then HbpR_CBP6_ should also be able to recognize OH-PCBs and be used for indirect detection of PCBs.

In this study, the HbpR_CBP6_ sensing module and the *bphAB* genes (encoding BphA and BphB enzymes that convert PCBs to OH-PCBs) derived from the *B. xenovorans* were integrated into genetically clear *E. coli* cells, and a highly sensitive WCB for the detection of PCBs was successfully constructed. After optimization, the best WCB could detect various PCB homologues, including 2-CBP, 4-CBP, 3-CBP 2,3-diCBP, 2,2’-diCBP, with detection limits ranging from 0.06 μM to 1 μM. Moreover, to provide a user-friendly solution for daily PCBs analysis, the optimized sensor cells were finally embedded in a gelatin-based hydrogel to prepare a smartphone-based detection program, thereby facilitating a smart, economical, real-time and on-site PCB monitoring in the real-world.

## 2. Materials and methods

### 2.1 Strains and chemicals

*E. coli* strains DH5α, TOP10, DB3.1, BL21(DE3), and Rosetta (DE3) were purchased from Tsingke Biotech (Beijing, China). *E. coli* HUBU is a DH5α-derived mutant strain generated in our laboratory (He et al., 2024). Analytical grade inducers, including 2-hydroxybiphenyl (2-HBP), 2-chlorinated biphenyl (2-CBP), 3-chlorinated biphenyl (3-CBP), 4-chlorinated biphenyl (4-CBP), 2,3-dichlorinated biphenyl (2,3-diCBP), 2,2’-dichlorinated biphenyl (2,2’-diCBP), and 4,4’-dichlorinated biphenyl (4,4’-diCBP), were purchased from Sigma-Aldrich. PCR-related enzymes were purchased from Vazyme Biotech (Nanjing, China). Genes and primers (**Table S1**) were synthesized by Sangon Biotech (Shanghai, China). Tryptone and yeast extract for Luria broth (LB) medium were purchased from Oxoid (Hampshire, UK).

### 2.2 Construction of plasmids and whole-cell biosensors

Plasmid construction was carried out using standard molecular biology techniques to create the sensing plasmids. The genes encoding HbpR_wt_ or HbpR_CBP6_ protein and the EGFP reporter were connected to the ends of P_hbpR_ and P_hbpC_ promoter by fusion PCR, respectively, to form an HbpR_wt_ or CBP6-P_hbpR_-P_hbpC_-EGFP expression cassette, which was then inserted into the *Sal* I/*Eco*R I dual-digested pUC57 vector to obtain the final plasmids pUC57-HbpR_wt_-P_hbpC_-egfp and pUC57-HbpR_CBP6_-P_hbpC_-egfp, respectively. To generate a PCB degrading plasmid, the genes encoding the BphAB enzymes were placed downstream of the strong constitutive promoter PUTR to create a sensor gene expression cassette (Zhou et al., 2021). This cassette was then cloned into the *Bam*H I/*Hin*d III dual-digested pMD18-T vector to generate the final plasmid pMD18-T-PUTR-bphAB.

The sensor plasmids pUC57-HbpR_wt_-P_hbpC_-egfp and pUC57-HbpR_CBP6_-P_hbpC_-egfp were transformed into different *E. coli* strain to obtain the first-stage WCBs, respectively. Similarly, seven second-stage WCBs were obtained by transforming the sensor plasmid pUC57-HbpR_CBP6_-P_hbpC_-*egfp* and the PCB degrading plasmid pMD18-T-P_UTR_-*bphAB* into different *E. coli* strains, respectively.

### 2.3 Cell growth, fluorescence assays and data analysis

For each whole-cell biosensor, 3-4 single clones were inoculated into 100 mL LB medium supplemented with antibiotic (kanamycin 50 μg/mL and ampicillin 100 μg/mL), and incubated at 37°C and 220 rpm. Once an optical density at 600 nm (OD_600_) of 0.6-0.8 was reached, the cell pellets were harvested by centrifugation at 4°C and 3800 rpm for 10 minutes, followed by washing three times with 3-(N-morpholino) propane sulfonic acid buffer containing 0.2% glucose (MOPS-G) and resuspended in MOPS-G buffer to an OD_600_ of 0.6. Next, 4 mL of suspended cells were treated with various PCB inducers at a ratio of 1000:1 to achieve final concentrations ranging from 0.01 to 1000 μM and then incubated at 30°C and 180 rpm for 6 hours. Afterwards, 200 μL of the induced sample was transferred to a microplate reader, and the fluorescence intensity and OD_600_ were measured using a fluorescent microplate reader (Spectra M2, Molecular Devices). The fluorescence value of each sample was calculated by normalizing fluorescence intensity to OD_600_. Finally, the resulting data were analyzed and plotted using GraphPad Prism software.

### 2.4 Structural analysis

The primary amino acids sequence of HbpR_wt_ protein was input into the Alphafold 2, an open-source online prediction website in Google Colab, to generate a simulated structure. Using this as a template, we then simulated the structure of the HbpR_CBP6_ variant using YASARA software. According to the chemical formula of the inducer, the 3D structure of each inducer was formed using ChemDraw software, followed by energy minimization using YASARA software to obtain the optimized structure. Afterwards, the inducer structure was docked with the 3D structure of HbpR_wt_ or HbpR_CBP6_ using YASARA software, and the most favorable position and orientations were identified using standard procedures; finally, Discovery Studio software was used to analyze the interaction forces in the HbpR protein-substrate complex structures.

### 2.5 Immobilization of whole-cell biosensors in gelatin hydrogels

To immobilize whole-cell biosensor in hydrogels, we first covered a glass slide with 1 mm thick tape, and the punched a 7 mm diameter hole in the tape using an infrared laser perforator. Next, we cultivated the biosensor *E. coli* BL21(DE3)/HbpR_CBP6_-bphAB in LB medium supplemented with kanamycin and ampicillin until the cell OD_600_ reached 0.6-0.8. The cells were then harvested by centrifugation at 4000 rpm for 2 minutes and washed 2-3 times with MOPS-G buffer. We then resuspended the cell pellet to an OD_600_=10 by MOPS-G buffer containing 16 U/mL transglutaminase and 10% gelatin to obtain a homogeneous mixture. After these steps, 15 μL of the homogenized mixture was pipetted into the wells on the glass slide and the gel was cross-linked at 37 ℃ for 20 minutes. The cross-linked gelatin cell mixture was mixed with different concentrations of 2-CBP and induced for 6 hours. The glass slide was then photographed under a blue light detector and the fluorescence intensity was quantified using ImageJ software.

## 3. Results and Discussions

3.1 Construction of transcriptional factor HbpR_CBP6_-based *E. coli* whole-cell biosensor

Two HbpR/P_hbpC_ sensing circuits were reported to function as whole-cell biosensors (WCBs) for the detection of 2-HBP and 2-CBP (Beggah et al., 2008). Therefore, we reconstructed these two WCBs, DH5α/HbpR_wt_ and DH5α/HbpR_CBP6_, by introducing the HbpR sensing circuits into *E. coli* DH5α cells. Similar to the report (Beggah et al., 2008), the WCB based on the HbpR_CBP6_ transcriptional factor variants were significantly more sensitive to the substrate 2-HBP than the DH5α/HbpR_wt_ WCB (**Fig. 1**). For the 2-CBP target, only the DH5α/HbpR_CBP6_ biosensor responded at concentrations above 75 μM, with a signal-to-noise (SNR) of 3.66. This is obviously worse than the result reported by Siham et al. (around 1.5 μM). We speculate that this may be due to the inevitable differences in experimental conditions and detection equipment (flow cytometry vs. microplate reader). Overall, the current WCB still need to be improved.

**Figure 1.**
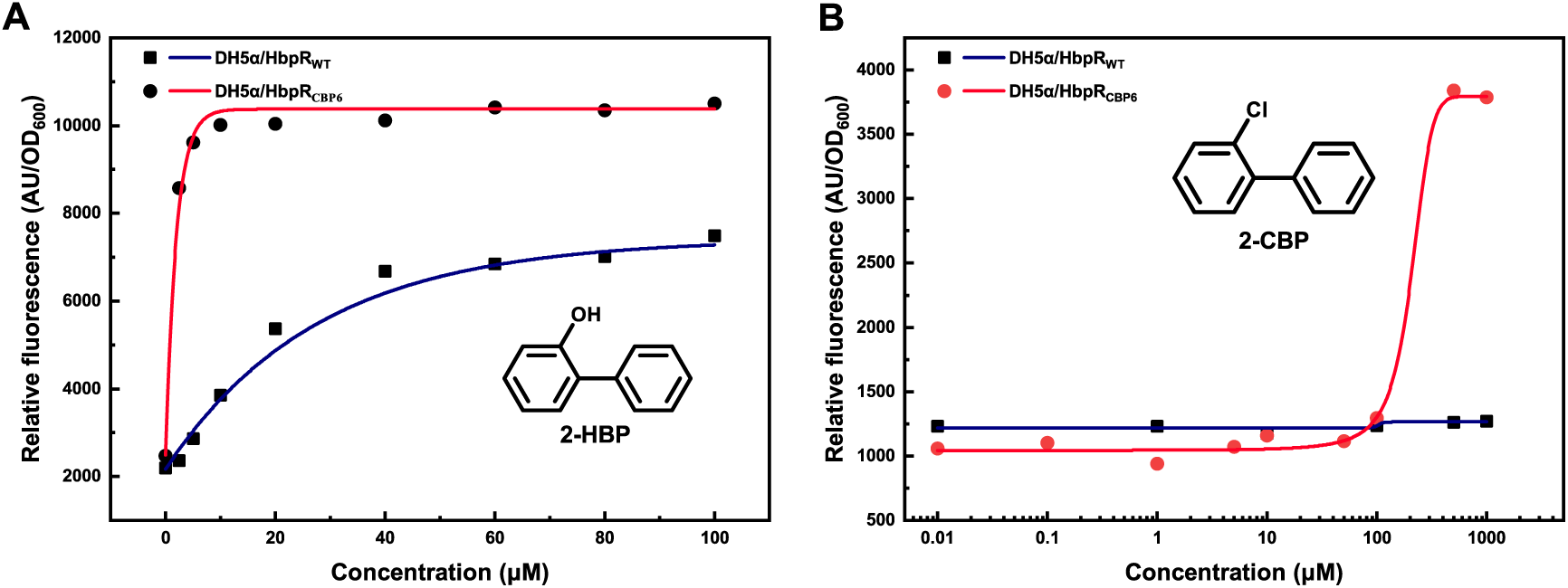
HbpR- based whole-cell biosensor for detection of 2-HBP and 2-CBP. Dose-response curves for wild-type (DH5α/HbpR_wt_) and variant (DH5α/HbpR_CBP6_) whole-cell biosensor used to detect 2-HBP (A) and 2-CBP (B) after induction for 6 hours.

### 3.2 Chassis genotype is critical for whole-cell biosensors

The genotype of chassis strains can profoundly affect the performance of the sensing pathways carried (Yagur-Kroll and Belkin, 2014). For example, Julien et al. observed a 10-fold increase in sensitivity of nickel-iron WCB when *E. coli* TD2158 was used as a chassis instead of *E. coli* K12 (Cayron et al., 2017). Our recent study also confirmed that the genotypes of chassis strain does have a significant impact on the effectiveness of WCB (He et al., 2024; Xian et al., 2024). Therefore, seven *E. coli* strains are selected as chassis carrying the *hbpR_CBP6_-P_hbpC_-eGFP* sensing circuit. These include three commonly used clonal strains (DH5α, Top10, DB3.1), two expression strains (BL21(DE3) and Rosetta (DE3)), and a mutant strain (HUBU) derived from DH5α strain, which served as the best chassis for hypersensitive salicylic acid WCB (He et al., 2024).

We observed that biosensors carried by BL21(DE3), Rosetta (DE3), and HUBU strains show significantly enhanced signal intensity and lower LOD compared to the DH5α strain. In contrast, the biosensor-carrying Top10 strain displays a worse response than DH5α, and biosensor-carrying DB3.1 did not respond to 2-CBP at all. Among these strains, the best chassis BL21(DE3) and Rosetta (DE3) show approximately three times lower LOD (from 75 μM to 23.5 μM) and 2.7-fold increased signal-to-noise ratio (SNR) than the DH5α strain (**Fig.2**).

**Figure 2.**
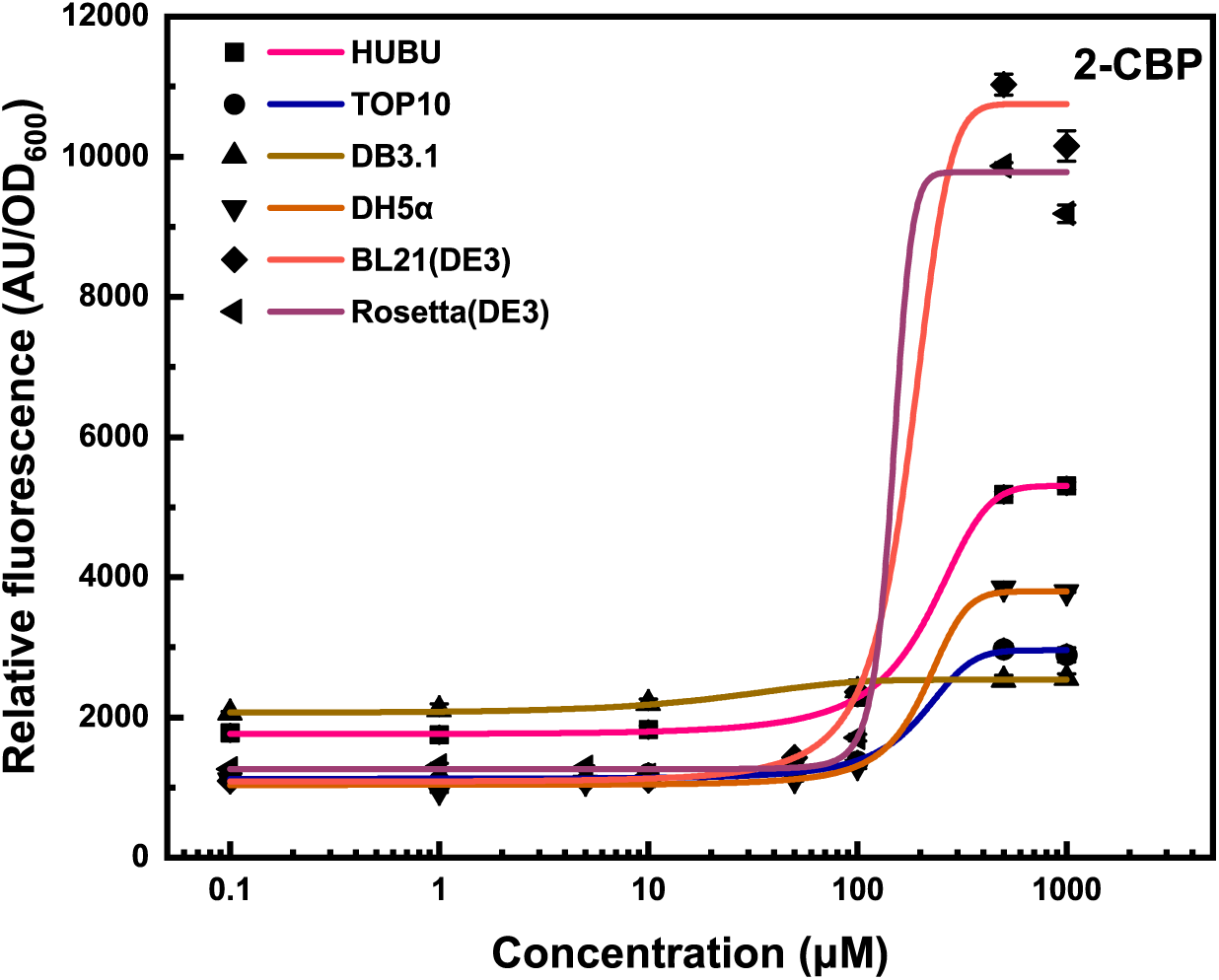
Dose-response curves of HbpR_CBP6_-based whole-cell biosensor with different *E. coli* chassis for detection of 2-CBP.

In this study, we found that the most effective chassis BL21(DE3) and Rosetta (DE3) were both of expression type. This observation is similar to another study in which expression strains BL21(DE3) and MG1655(DE3) were identified as optimal hosts for chlorobenzoic acids (CBA) whole-cell biosensor, whereas clonal strains DH5α, Top10, DB3.1 were found to be relatively poor chassis (Xian et al., 2024). Through genotype comparison of these strains, we find that all positive strains, including BL21(DE3), Rosetta (DE3), and MG1655(DE3), contained functional *recA* and *endA* genes in their genomes. In contrast, most of the poorer chassis were *recA*-*endA-*deficient strains. The *recA* and *endA* genes encode DNA recombinase and endonuclease, respectively, which appear to play major roles in DNA polymerization and recombination (Bukhamsin et al., 2022; Sun et al., 2014). We hypothesize that these enzymes may facilitate greater accessibility of cis-acting elements (promoters) in plasmid DNA to trans-acting regulatory proteins (e.g., XylS or HbpR_CBP6_), thereby enhancing transcription of downstream reporter genes. Therefore, WCBs based on these expression-type strains display higher signal output than those strains lacking the *recA1-endA1* genes, such as DH5α, Top10, DB3.1 strains. This hypothesis is further supported by studies on DH5α-related HUBU strains, in which the *recA1*(D161G) mutation in the HUBU strain may allow DH5α to regain the recombination function lost due to *recA1* gene mutation, resulting in the regeneration of recombination activity of later sensing strains (He et al., 2024). Finally, the HUBU strain carried a salicylic acid (SA) biosensor, and our PCB biosensors all show higher sensitivity and signal strength than the DH5α strain. Poor chassis, on the other hand, contain various genetic defects in energy or amino acid synthesis pathways. For example, DH5α strain lacks the GTP-pyrophosphate kinase synthesis gene *relA*, Top10 is a strain defective in the leucine synthesis gene, and DB3.1 lacks the γ-glutamyl phosphate reductase gene *proA2* (Durfee et al., 2008; Jeong et al., 2017; Kwon et al., 2020). These defects may slow down the expression of sensing proteins and reporter genes, resulting in poorer WCB chassis.

### 3.3 Developing PCB WCBs by additionally introducing *bphAB* degradation pathway

Since HbpR_CBP6_ has a higher affinity for OH-PCBs, this inspired us to convert PCB into a higher affinity OH-PCB intermediate for indirect detection of PCB. To achieve this, we integrated the *bphAB* genetic circuit derived from the *B. xenovorans* LB400 strain (Hofer et al., 1994) into the BL21(DE3)/HbpR_CBP6_ WCB to generate a novel BL21(DE3)/HbpR_CBP6_-*bphAB* WCB. These two genes encode *bphA* and *bphB* enzymes that convert PCB to OH-PCB, which may serve as a higher affinity ligand for the HbpR_CBP6_ transcriptional factor. Our results show that the additive *bphAB* genetic circuit does not affect the signal intensity of the BL21(DE3)/HbpR_CBP6_-*bphAB* WCB. Furthermore, the detection limit of 2-CBP was significantly reduced by about 68.8-fold (from 23.5 μM to 0.34 μM). This is the lowest LOD achieved for 2-CBP to date and is 4.4-fold lower than the biosensor (1.5 μM) reported by Siham Beggah et al (**Fig. 3**).

**Figure 3.**
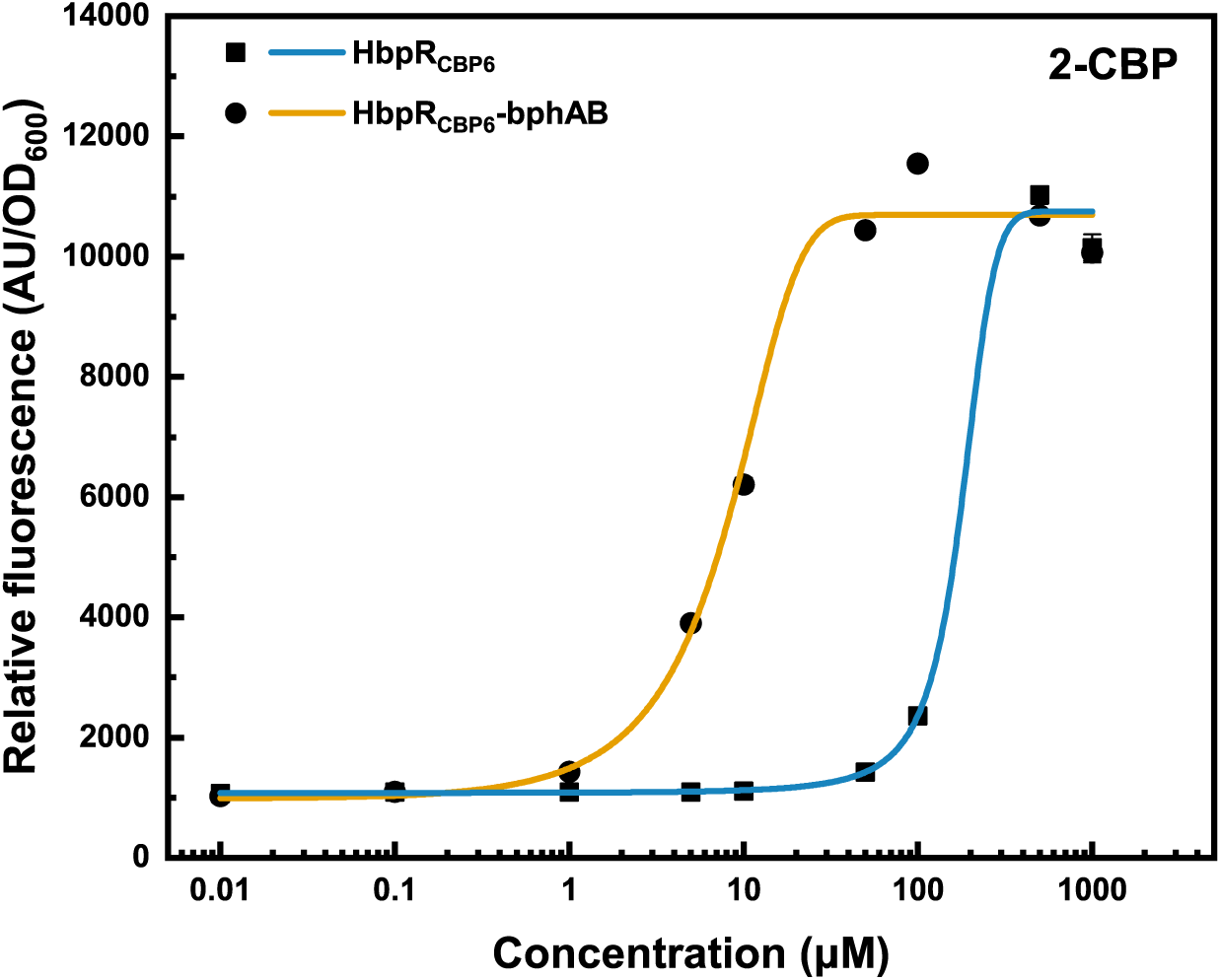
Dose-response curves of HbpR_CBP6_- and *bphAB-* based whole-cell biosensor with *E. coli* BL21 (DE3) for detection of 2-CBP.

**Figure 4.**
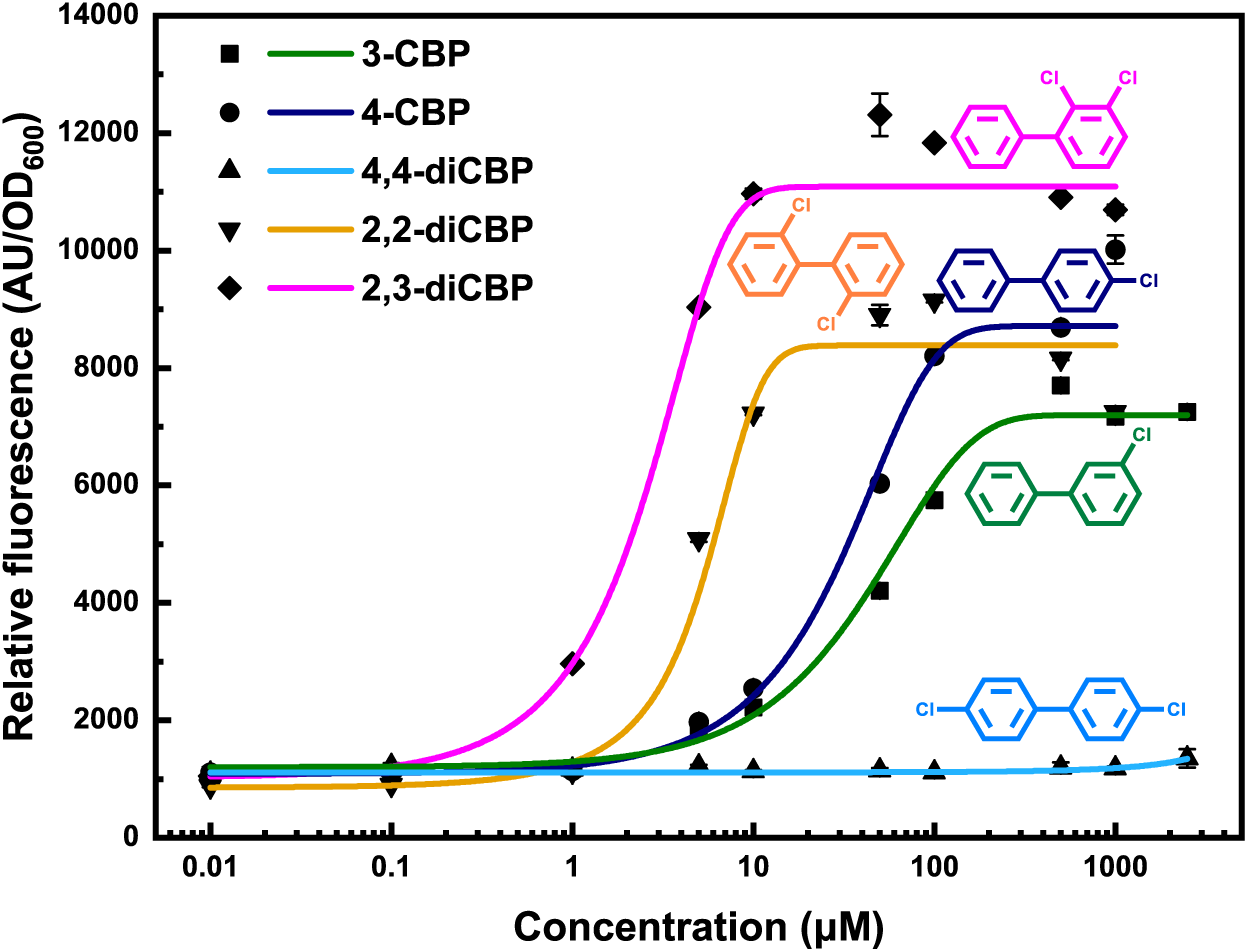
PCB detection range of the BL21(DE3)/HbpR_CBP6_-*bphAB* whole-cell biosensor. 3-CBP: 3-chlorinated biphenyl; 4-CBP: 4-chlorinated biphenyl; 4, 4’- DiCBP: 4, 4’-dichlorinated biphenyl; 2, 2’-DiCBP: 2, 2’-dichlorinated biphenyl; 2, 3- DiCBP: 2, 3-dichlorinated biphenyl.

The increased sensitivity of BL21(DE3)/HbpR_CBP6_-*bphAB* WCB can be attributed to the catalytic conversion of the lower affinity target 2-CBP to the higher affinity hydroxylated 2-CBP (2,3-diOH-2-CBP) target catalyzed by *bphA-bphB* enzymes. This speculation is further supported by the enhanced molecular interaction (**Fig. 5**) and lower binding energy (**Fig. 6**) of HbpR_CBP6_ on 2,3-diOH-2-CBP than 2-CBP. Prior to our study, there have been sporadic reports focusing on monitoring PCB metabolic intermediates for indirect PCB detections. These biosensors are typically developed in *Pseudomonas* species, which have their own pathways for the degradation of PCB, thereby there is potential interference from autocrine pigments or unknown metabolic pathways (Gavlasova et al., 2008; Liu et al., 2010). Contrary to these reports, our WCB is the first example of transplanting a heterogenous PCB metabolic pathway into an *E. coli* species for indirect PCB detections. *E. coli* strains have become popular chassis for synthetic biology due to their clear genetic background and systematic operational procedures. These advantages allow us to develop highly reactive, low background BL21(DE3)/HbpR_CBP6_-*bphAB* WCB through chassis screening and genetic reorganization, and could also promote the practical application of PCB WCBs in academia and industry.

**Figure 5.**
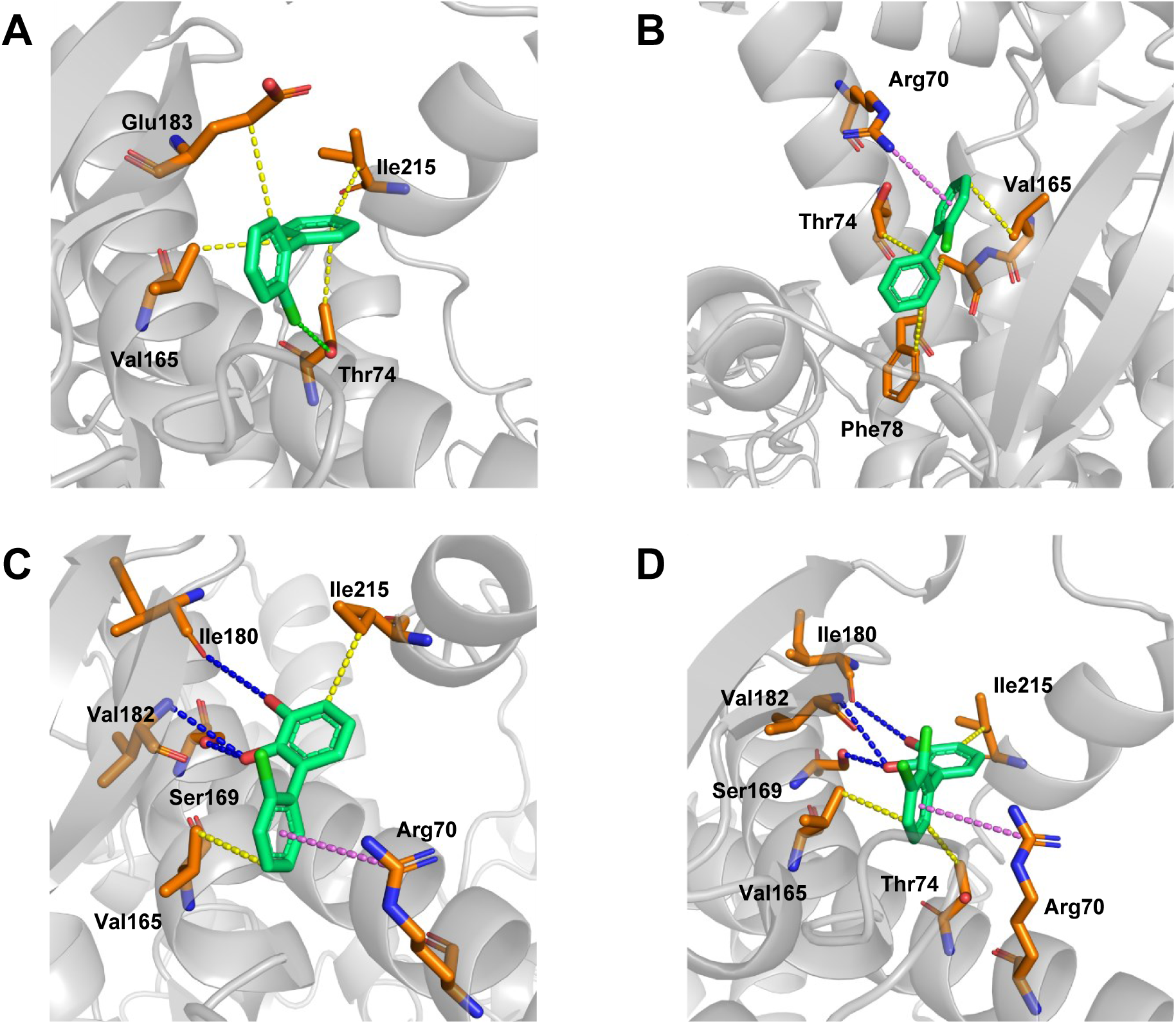
Interaction forces between HbpR and 2-CBP or its analogs. (A) HbpR_wt_ and 2-CBP. (B) HbpR_CBP6_ and 2-CBP. (C) HbpR_CBP6_ and 2, 3-diOH-2-CBP. (D) HbpR_CBP6_ and 2, 3-diOH-2, 3-diCBP. Yellow indicates hydrophobic interactions, pink indicates π-cation interactions, blue indicates hydrogen bond interactions, and green indicates halogen bond.

**Figure 6.**
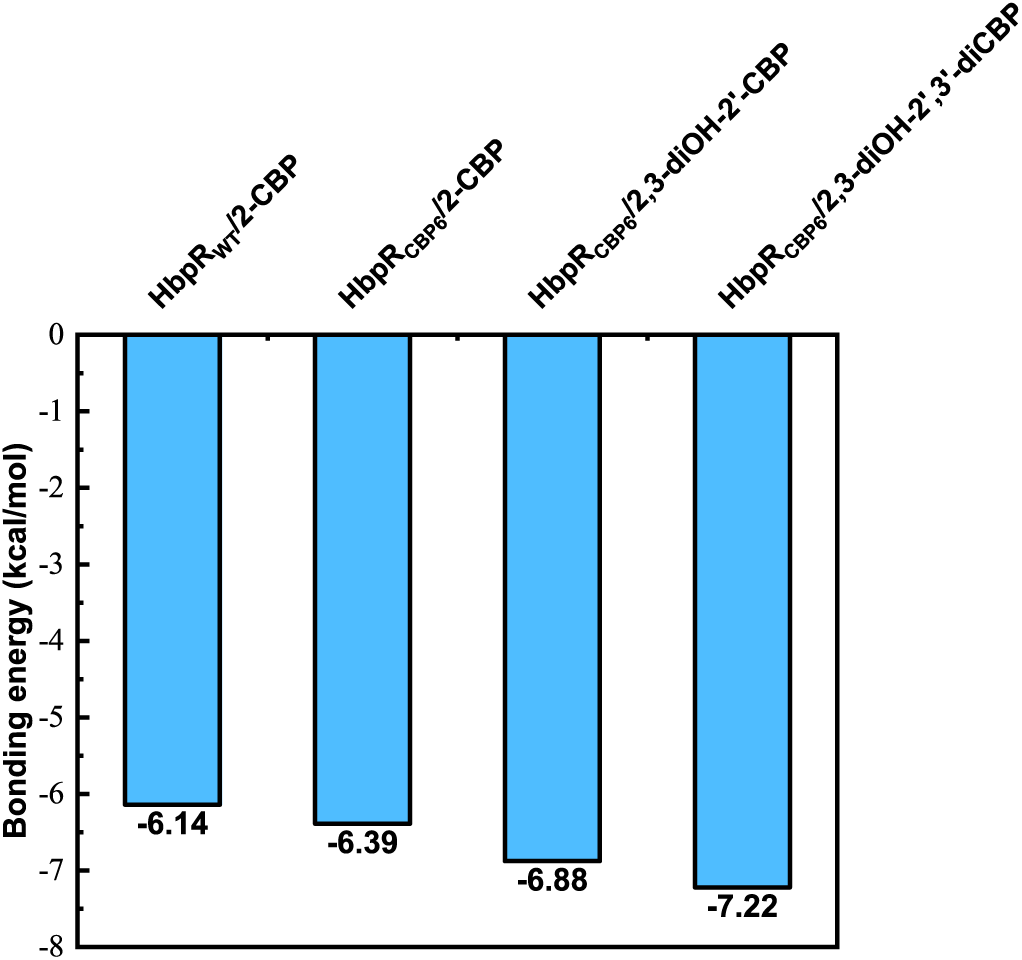
Bonding energies of HbpR_wt_ with 2-CBP, HbpR_CBP6_ with 2-CBP, 2, 3-diOH-2-CBP, and 2, 3-diOH-2, 3-diCBP, respectively.

### 3.4 Detection spectrum of BL21(DE3)/HbpR_CBP6_**-***bphAB* whole-cell biosensor

PCBs contain a variety of homologues with different numbers of chlorine atoms and substitution positions. This is a critical issue for instrumental and ELISA based analysis, as they generally prefer single targets. Whole-cell biosensors capable of detecting multiple homologues simultaneously would offer tremendous advantageous for real-world PCB detection. We have demonstrated that the optimized BL21(DE3)/HbpR_CBP6_-*bphAB* WCB was highly responsive to 2-CBP (**Fig. 3**). To evaluate its potential for detecting complex PCBs, we further evaluated its ability to respond against 5 other PCB homologues, including 3-chlorobiphenyl (3-CBP), 4-chlorobiphenyl (4-CBP), 2,3-dichlorobiphenyl (2,3-diCBP), 2,2’-dichlorobiphenyl (2,2’-diCBP), and 4,4’- dichlorobiphenyl (4,4’-diCBP). Our results show that the final WCB displays sensitive responses to 4 test targets, including 2,3-diCBP, 2,2’-diCBP, 4-CBP and 3-CBP, with LODs close to 0.06 μM, 0.3 μM, 0.88 μM and 1 μM, respectively (**Fig. 4**). However, no response to 4,4’-diCBP was observed.

The highest and lowest responses of the BL21(DE3)/HbpR_CBP6_-bphAB biosensor to the 2,2’- diCBP and 4,4’-diCBP targets, respectively, may be due to the substrate recognition of different PCB substrates by the BphAB enzymes and determined through different regiospecificity. Indeed, the *B. xenovorans* LB400-derived BphA enzyme is a 3,4-dioxygenase that typically introduces hydroxyl groups into the positions 3 and 4 of the phenyl ring in PCBs. Therefore, the 2,2’-diCBP product catalyzed by BphA is mainly composed of 2,3-dihydro-2,3-dihydroxy-2’-chlorobiphenyl and 3,4- dihydro-3,4-dihydroxy-2,2’-dichlorobiphenyl. These compounds were subsequently dehydrogenated to 2,3-dihydroxy-2’-chlorobiphenyl (2,3-diOH-2-CBP) and 3,4-dihydroxy-2,2’- dichlorobiphenyl (3,4-diOH-2,2’-CBP) under the catalysis of BphB dehydrogenase, of which the former is the main product (Li et al., 2020). It is worth noting that 2,3-diOH-2-CBP is also the hydroxylation product of 2-CBP catalyzed by bphAB enzymes and can be well recognized by the HbpR_CBP6_ transcriptional factor. Therefore, the BL21(DE3)/HbpR_CBP6_-*bphAB* WCB exhibited similar LOD and SNR values to the 2,2’-diCBP and 2-CBP targets.

In contrast, the reactivity of BL21(DE3)/HbpR_CBP6_-*bphAB* WCB towards 4,4’-diCBP was negligible, which may be due to the lower selectivity of the BphA enzyme for the para-substituted PCBs substrates (ortho > meta > para) (Gomez Gil, 2006). This preference made it difficult for the BphA enzyme to recognize 4,4’-diCBP and convert it into a hydroxylated intermediate for downstream HbpR_CBP6_ detection. As a result, negligible reactivity was obtained with 4,4’-diCBP even at a concentration of 1000 μM (**Fig. 4**). Abundant enzyme resources and rapid development of enzyme directed evolution technology have greatly promoted the development of BphA enzymes with expanded substrate spectrum. We anticipated that more desired variants will be introduced into sensitive WCBs to enable broader PCBs detection.

### 3.5 The recognition mechanism PCB and OH-PCB by HbpR_CBP6_

To date, a major challenge in developing highly sensitive WCBs for PCB detection has been the lack of efficient PCB sensing proteins. As a pioneer in direct recognition of 2-CBP, HbpR_CBP6_ provides valuable insights for de novo design or evolution of artificial PCB-sensing proteins. Therefore, structural simulations and molecular docking were carried out to generate complexes of HbpR protein with PCBs and PCB analogs. From these models, we found that the interaction of wild-type HbpR with 2-CBP primarily relied on weak hydrophobic interactions and halogen bonds (Brammer et al., 2001) (**Fig. 5A**), whereas the HbpR_CBP6_ variant mainly used hydrophobic interactions and residue Arg70 to participate in the well-known and important π-cation interactions with 2-CBP (**Fig. 5B**). π-cation interaction has been recognized as strong noncovalent forces and shown to affect protein-protein and protein-small molecule interactions (Dougherty, 2007; Gokel et al., 2001; Torrice et al., 2009). Therefore, we hypothesize that the expanded recognition of 2-CBP by HbpR_CBP6_ protein variants is due to their stronger interactions, thereby enhancing the binding energy between HbpR_CBP6_ protein and the substrate 2-CBP, which may enable the target to induce conformational changes in the HbpR_CBP6_ transcriptional factor to promote RNA polymerase binding and gene transcription. This hypothesis is further supported by the higher binding energy between the HbpR_CBP6_ variant and 2-CBP (-6.39 kcal/mol) than its wild-type counterpart (-6.14 kcal/mol) (**Fig. 6**).

Structural analysis between HbpR_CBP6_ and the hydroxylated 2-CBP (2,3-diOH-2-CBP) and 2,3- diCBP (2,3-diOH-2,3-diCBP) were also carried out to further verify the speculation that the improved sensitivity of BL21(DE3)/HbpR_CBP6_-*bphAB* WCB for PCB detection was facilitated by the *bphAB* enzymes-mediated conversion of PCBs into higher-affinity OH-PCBs. Compared with its targets of 2-CBP and 2,3-diCBP, HbpR_CBP6_ appear to form a more complex interaction network with its hydroxylated ligands, including weak hydrophobic interactions, strong π-cation interactions, and hydrogen bond interactions between hydroxyl groups of hydroxylated intermediates and surrounding residues such as Ser169, Ile180, and Val182 (**Fig. 5C, D**). These interactions obviously enhanced the binding affinity between HbpR_CBP6_ and hydroxylated 2-CBP and 2,3-diCBP, thereby promoting the indirect detection of PCBs. Likewise, this observation can be further supported by the lower binding energies of HbpR_CBP6_ with 2,3-diOH-2-CBP and 2,3-diOH-2,3-diCBP than 2-CBP and 2,3-diCBP (**Fig. 6**).

With the increasing public concern about health issues, the demand for affordable and user-friendly sensing devices has grown significantly. Smartphones are almost one-to-one devices, offering a promising medium for diagnose and biosensing (Yang et al., 2023). Using customized Apps, smartphones can quickly analyze target samples by capturing and digitizing signals (colorimetric, fluorescent, electrochemical, and surface plasmon resonance (SPR)) (Vashist et al., 2015). Here, we aimed to integrate our laboratory BL21(DE3)/HbpR_CBP6_-*bphAB* WCB with a smartphone application. To achieve this, we fist immobilized WCB cells into a hydrogel matrix via transglutaminase-gelatin cross-linking and evaluated their sensing capability by exposing them to different concentrations of 2-CBP. Analysis using Image J software shows that a good linear relationship between the logarithm of 2-CBP concentrations and fluorescence intensities in the range of 1 μM to 100 μM (**Fig. 7A, C**), indicating that the immobilized hydrogel WCB has good sensing performance.

**Figure 7.**
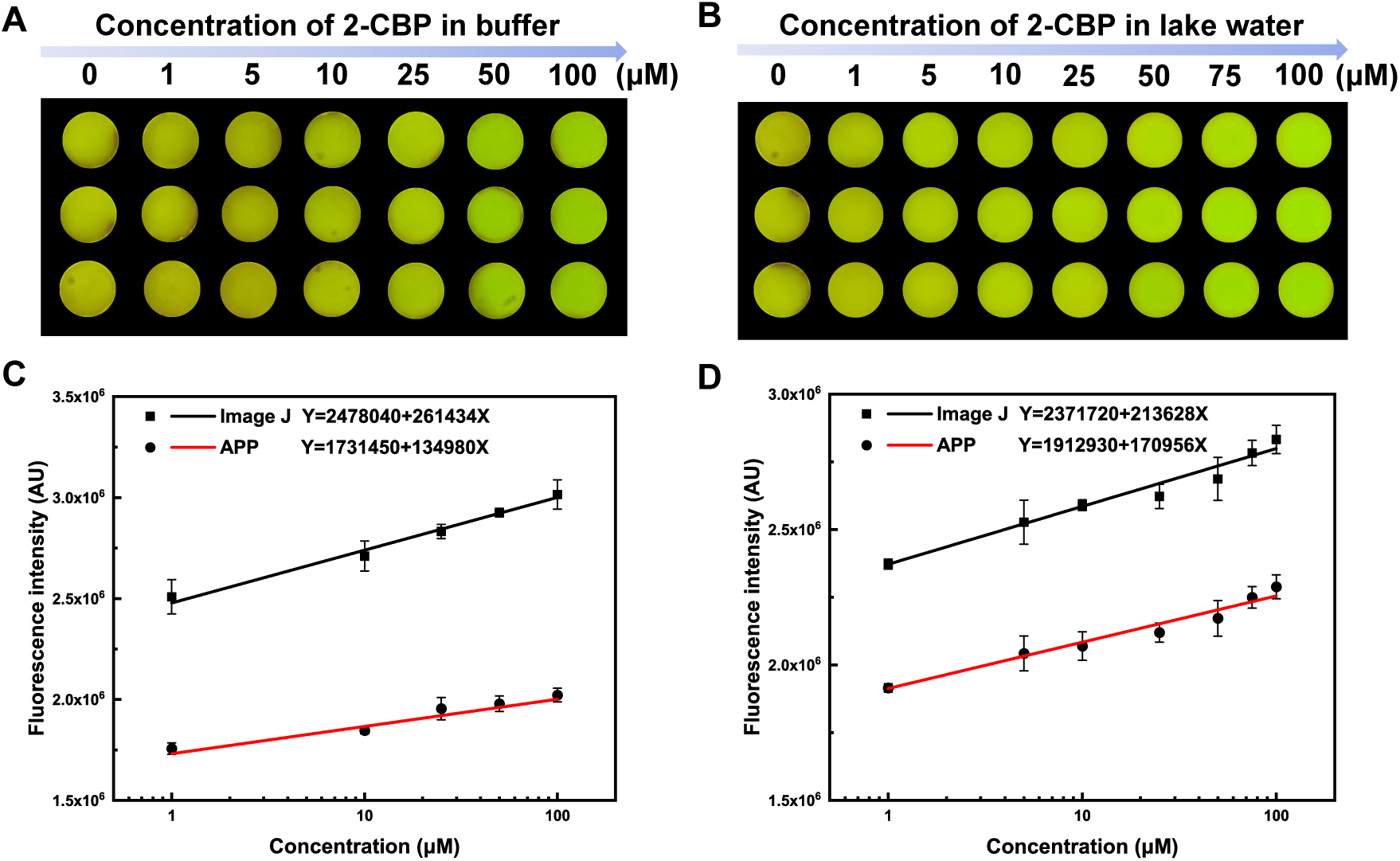
Detection of 2-CBP with hydrogel biosensor. (A) Fluorescence changes of 2- CBP at different concentration in MOPS buffer; (B) Fluorescence changes of 2-CBP at different concentration in lake water; (C) Linear relationship between smartphone APP detection of 2-CBP in MOPS buffer; (D) Linear relationship between smartphone APP detection of 2-CBP in lake water.

Subsequently, following a similar smart procedure outlined in our recent publication (He et al., 2024), we developed a portable device to capture PCBs-induced fluorescence of immobilized hydrogel biosensors using a smartphone camera, then digitized and output it as a specific image. We evaluated the performance of this smart program by analyzing the above 2-CBP standard solutions. The data output from smartphone exhibited a robust linear function relationship with 2-CBP concentration, with accuracy comparable to Image J software (**Fig. 7C**). We then tried to apply this detection procedure to analyze real environmental samples, but such samples were inconvenient to collect. Therefore, we instead prepared spiked environmental samples by adding 2-CBP to urban lake water. As expected, this smart procedure accurately reflected the spiked samples in the range of 1 μM to 100 μM (**Fig. 7B, D**), almost no interference from other substances in lake water. Overall, these investigations demonstrate the huge potential of our smartphone-based detection devices for intelligent, cost-effective, and rapid analysis of PCBs in the real-world.

## 4. Conclusion

In summary, we have engineered a sensitive *E. coli* whole-cell biosensor BL21(DE3)/HbpR_CBP6_-*bphAB* by transplanting the *bphAB* genes derived from the *B. xenovorans* LB400 strain into *E. coli* cells and conducted comprehensive strain screening to detect various PCBs in a highly sensitive manner. This WCB exhibits sensitive and broad responsiveness to various PCB homologues, including 2,3-diCBP, 2,2’-diCBP/2-CBP, 4-CBP/3-CBP, with a detection limit ranging from 0.06 μM to 1 μM. We then devised a user-friendly detection procedure by integrating immobilized hydrogel WCB cells, portable imaging device, and a customized smartphone App to facilitate a smart, cost-effective, and rapid detection of PCBs in environmental samples. We believe these studies will promote biomonitoring and bioremediation of PCBs contaminants, thereby contributing to the protection of the environment and human health.

## Acknowledgements

This work was supported by National Key R&D Program of China (No.2018YFA0901100), National Natural Science Foundation of China (32370101) and Fundamental Research Funds for the Central Universities (buctrc202131).

