## Supplemental Table 1 for "A smartphone-based genetically recombinant whole-cell biosensor for highly sensitive monitoring of polychlorinated biphenyls (PCBs)"

Table S1 Primer sequences used in the experiment

| Primer | Sequences |
| --- | --- |
| TER-F | GCCTCTGCAGTCGACGACTCTCTAGCTTGAGGCATCAAATAAAACGAAAGGCTCAG |
| H1-R | CCCTCCTTATAAAGTTAATCTTTAGTTAGTTAGGGCGGCCGCTGTTATCACGTCTCCGC |
| H2-F | TAACTTTATAAGGAGGGAAAAACATATGGTGAGCAAGGGC |
| TER-R | ACGGCCAGTGAATTCATTCTCACCAATAAAAAACG |
| I101V-F | AAGACACCACCTGCATAAACCTGTTCTTTGATAGAGGATATACCTTG |
| I101V-R | TATCAAAGAACAGGTTTATGCAGGTGGTGTCTTGCACG |
| D128N-F | TCTTCATATTGACCGCATTTAGCGCCGATGATC |
| D128N-R | GCTAAATGCGGTCAATATGAAGAGCATGGATTATCATGCTGAGGC |
| Bph-F | CCGGGGATCCTCTAGAGATATTGGCTATCACATCCGACAC |
| Bph-R | AAACGACGGCCAGTGCCAAGCTTGCATGCCTGCAGGTCGACGATTACTGTCTCC |
